# Modeling Zika Virus Congenital Eye Disease: Differential Susceptibility of Fetal Retinal Progenitor Cells and iPSC-Derived Retinal Stem Cells to Zika Virus Infection

**DOI:** 10.1101/128405

**Authors:** Deisy Contreras, Melissa Kaye Jones, Laura E. Martinez, Vineela Gangalapudi, Jie Tang, Ying Wu, Jiagang J. Zhao, Zhaohui Chen, Shaomei Wang, Vaithilingaraja Arumugaswami

## Abstract

Zika virus (ZIKV) causes microcephaly and congenital eye disease that is characterized by macular pigment mottling, macular atrophy, and loss of foveal reflex. The cell and molecular basis of congenital ZIKV infection are not well understood. Here, we utilized a biologically relevant cell-based system on human fetal retinal pigment epithelial cells (FRPE) and iPSC-derived retinal stem cells (iRSCs) to model ZIKV-ocular cell injury processes. FRPEs were highly susceptible to ZIKV, resulting in apoptosis and decreased viability, whereas iRSCs showed reduced susceptibility. Transcriptomics and proteomics analyses of infected FRPE cells revealed the activation of innate immune and inflammatory response genes, and dysregulation of cell survival pathways, mitochondrial transmembrane potential, phagocytosis, and particle internalization. Nucleoside analogue drug treatment inhibited ZIKV replication and prevented apoptosis. In conclusion, ZIKV affects ocular cells of different developmental stages resulting in cellular injury and death, further providing molecular insight into the pathogenesis of congenital eye disease.

## Introduction

Zika virus (ZIKV) is a teratogenic vector-borne pathogen that causes detrimental congenital eye disease along with other congenital disorders, such as microcephaly, intrauterine growth restriction, and sensorineural hearing loss (Jampol and Goldstein, 2016). ZIKV is a member of the *Flaviviridae* family, comprised of other relevant human pathogens such as West Nile, Japanese Encephalitis, Dengue, and St. Louis Encephalitis viruses. ZIKV was first isolated from humans in 1952 in Uganda and Tanzania, and its main mode of transmission is vectorial through the bite of an infected *Aedes* species mosquito (*Aedes aegypti* or *Aedes albopictus*) (Hamel et al., 2016). However, new studies provide increasing evidence that ZIKV can be spread sexually from men or women, and through transplacental transmission in infected mothers (Kim and Shresta, 2016). ZIKV may gain access to the fetal eye through hematogenous route and direct ocular exposure to infected amniotic fluid *in utero* resulting in congenital eye disease. Approximately 80% of adults infected with ZIKV remain asymptomatic, while the remaining infected population exhibit mild symptoms, such as short-lived fever, rash, and joint pain, all of which are rarely life threatening (Hamel et al., 2016). ZIKV has also been linked to Guillain-Barré syndrome, which can damage nerve cells and cause temporary paralysis in infected adults (dos Santos et al., 2016; Lucchese and Kanduc, 2016; Parra et al., 2016).

Congenital ocular defects occur at a frequent rate worldwide and approximately 20 million children under the age of 18 years suffer from some type of congenital eye disorder, resulting in more than 1.4 million cases of blindness (Chen et al., 2009; Graw, 2003; Lupo et al., 2000). Human pathogens known to cause fetal and neonatal anomalies include *Toxoplasma gondii*, Rubella virus, Cytomegalovirus (CMV), and Herpes Simplex Virus 1 (HSV-1), referred to as TORCH infections, and can be acquired by intrauterine route (Kim and Shresta, 2016). A repertoire of clinical studies on ZIKV infected fetuses have found that infection leads to a spectrum of eye anomalies during the first or second trimester of pregnancy, such as macular pigment mottling, loss of foveal reflex, intra-retinal hemorrhages, chorioretinal atrophy, and blindness during the first or second trimester of pregnancy (de Paula Freitas et al., 2016; Ventura et al., 2016a; Ventura et al., 2016b; Vogel, 2016). Approximately 35% of microcephalic babies showed posterior pole pigmentary clumping and chorioretinal atrophy in the eyes (de Paula Freitas et al., 2016). ZIKV also causes structural eye anomalies such as microphthalmia and coloboma, cataracts, and intraocular calcifications in the eye (Rasmussen et al., 2016). However, the molecular mechanisms underlying the structural and developmental eye anomalies caused by ZIKV infection are not well understood.

Recently, immunocompromised mouse model systems have been utilized to reproduce key features of human ZIKV infection, including neuronal and ocular tissue tropism (Cugola et al., 2016; Lazear et al., 2016; Li et al., 2016; Miner et al., 2016). It has been shown that ZIKV has a broader cell tropism than other flaviviruses (Miner and Diamond, 2017). A study using an *ifnar1*^-/-^ knockout mouse model system demonstrated the presence of ZIKV in different parts of the eye, including the cornea, retinal epithelial cell layer, and tear fluid, which further contributed to conjunctivitis and uveitis (Miner et al., 2016). However, congenital eye disorder was not observed in *ifnar1*^+/-^ fetuses of C57BL/6 *ifnat1*^-/-^ infected mothers (Miner et al., 2016). A recent study reported that *in vitro* ZIKV infection of human retinal pigment epithelial (RPE) cells caused detrimental effects on membrane permeability by disrupting cell-to-cell junctions (Salinas et al., 2016). Altogether, fetal ocular infectious cell culture models are needed to study ZIKV pathogenesis during the early developmental stages of the eye.

Here, we utilized second-trimester human fetal retinal pigment epithelial (FRPE) progenitor cells and human induced pluripotent stem cell (iPSC)-derived retinal stem cells (iRSCs), analogous to first-trimester retinal progenitor cells, to recapitulate the pathogenic processes of congenital ZIKV infection *in vitro* by examining the initial ocular cellular response to infection and injury using the contemporary ZIKV clinical isolate PRVABC59. We found that ZIKV infection alters cell viability by inducing programmed cell death in FRPE cells. iRSCs show differential susceptibility to ZIKV infection. Global transcriptomics and proteomics analyses of FRPE cells revealed dysregulation of cell survival pathways and VEGF signaling, and activation of innate immune and inflammatory response genes during ZIKV infection. Further antiviral drug screening against ZIKV revealed that a nucleoside analogue, 6-Azauridine, inhibited viral replication and blocked ZIKV-mediated cell death in FRPE cells.

## Results

### Fetal retinal pigment epithelial cells are susceptible to ZIKV infection

To investigate the pathophysiology of ZIKV congenital eye infection, we employed an *in vitro* cell culture model system using human fetal retinal pigment epithelial cells (FRPE) from second-trimester of pregnancy to study their response to ZIKV infection and evaluate changes during ocular cell injury. For this study, we utilized a contemporary clinical ZIKV strain (PRVABC59), isolated in Puerto Rico during the 2015/2016 epidemic (Petersen et al., 2016). Fetal RPE cells were verified by the presence of retinal pigment epithelium-specific markers RPE65 and bestrophin (BEST1) (Figure S1). FRPE cells do not form well-defined tight junctions, as observed for adult RPE cells (Figure S1). The FRPE cells expressed several flaviviral cell entry receptors GRP76, SDC2, HSP90AB1, TYRO3, AXL, and MERTK (Figure S1B). To understand the ZIKV cytopathic effect (CPE) in human ocular cells, FRPE cells were infected with a multiplicity of infection (MOI) of 0.1, as previously described (Contreras and Arumugaswami, 2016). The FRPE cells showed active viral infection as evident by the presence of viral plaques on the cell monolayer (Figure 1A). The infected cells exhibited a distinctive raised pleomorphic phenotype and were detached from the cell’s monolayer (Figure 1A). ZIKV infection was confirmed by immunocytochemistry analysis of mock and infected FRPE cells using an antibody against the flavivirus envelope (Env) structural protein (Figure 1B). ZIKV genome replication was verified by RT-qPCR at 2 and 4 days post infection (dpi) (Figure 1C). To further characterize the cellular response to ZIKV infection, we examined cell viability by measuring the intracellular ATP content using Cell Titer-Glo assay. ZIKV infection of FRPE cells decreased cell viability at 2 and 4 dpi (20% vs. 60%) as compared to mock infected cells, which exhibited 100% viability (Figure 1D). We then examined the apoptotic effects of ZIKV on FRPE cells by measuring caspase 3/7 activity. At 4 dpi, an increase in caspase 3/7 activity was detected in ZIKV-infected cells, suggesting induction of programmed cell death (Figure 1E). FRPE cells were subjected to Annexin V staining to determine the early phase of apoptosis, as defined by viral plaques. At 3 dpi, FRPE cells show double staining for Annexin V and propidium iodide (PI), while mock infected cells show little or no measurable apoptosis (Figure S2). Overall, our data shows that the ZIKV-infected FRPE cells had increased caspase 3/7 activity and reduced cell viability due to robust viral infection.

**Figure 1.**
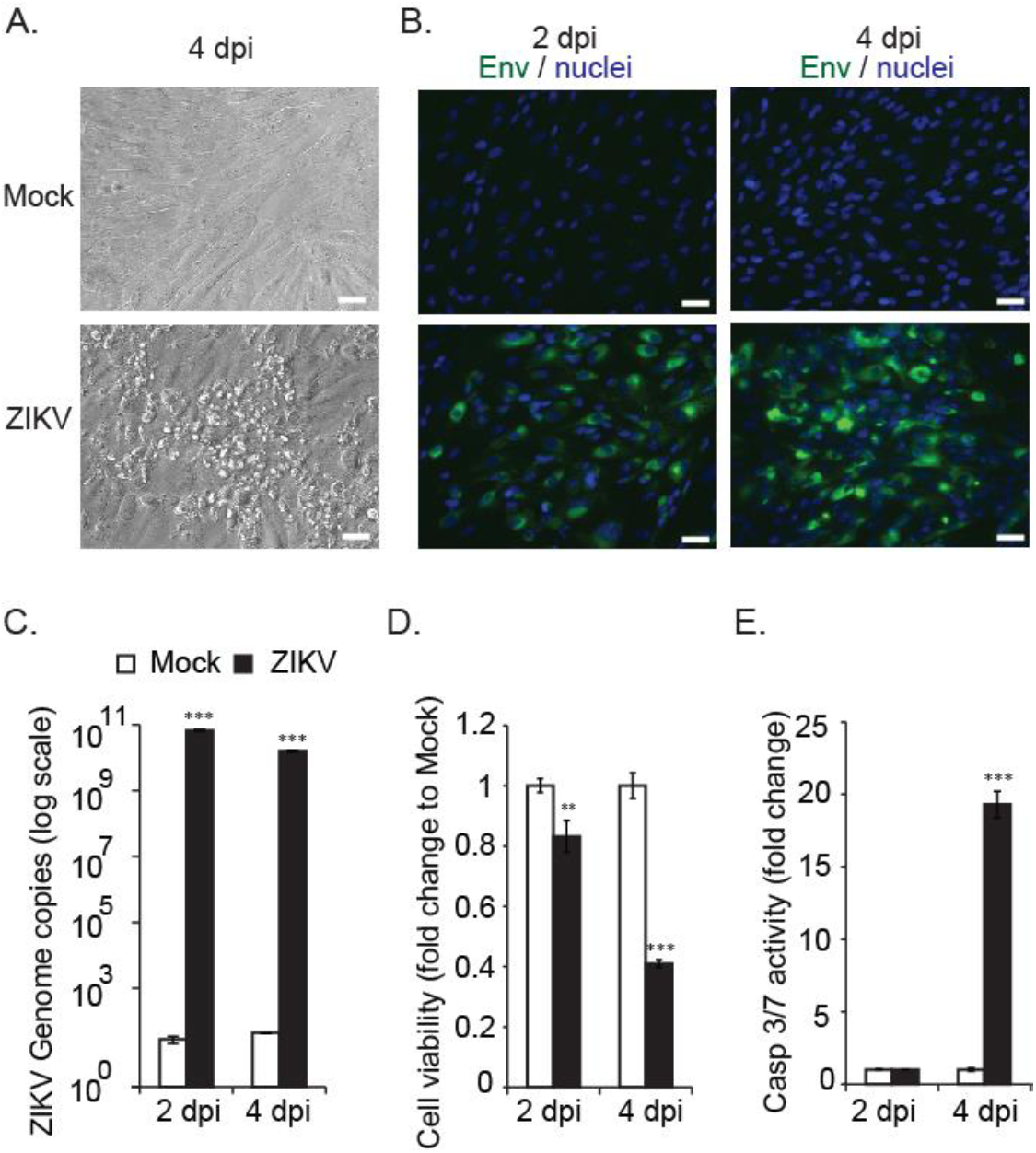
Fetal RPE cells are susceptible to ZIKV infection. (A) Bright field images of uninfected (mock) and ZIKV infected FRPE cells at 4 dpi. (B) Immunocytochemistry analysis of mock and ZIKV infected FRPE cells at 2 and 4 dpi. The presence of flavivirus envelope protein is shown in green and nuclei in blue. (C) Graph shows the ZIKV genome copies (log scale) of mock and ZIKV infected cells at the indicated timepoints. (D) Bar graph shows reduced cell viability for ZIKV infected cells. (E) Caspase 3/7 activity measured for mock and ZIKV-infected cells at 2 and 4 dpi. Scale bar: 25 μm. Student t-test was performed for statistical significance if the p value was p<0.05 (*); p<0.001 (**); and p<0.0001 (***). Representative data from three independent experiments are shown.

### Global Transcriptomics and Proteomics Analysis of ZIKV-infected fetal RPE cells

We then wanted to see the effects of ZIKV infection on FRPE cells at the molecular level by transcriptomics and proteomics analyses. RNA sequencing analysis of polyadenylated transcripts revealed that at 2 dpi, there were 2,040 differentially regulated genes upon ZIKV infection, with a *p*-value of less than 0.05 (Figure 2A and Table S2). At 3 dpi, there were 7,951 differentially regulated genes upon viral infection (Figure 2A and Table S3). Figure 2B provides a heat map of differentially expressed canonical interferon stimulated genes (ISGs) at 2 dpi (128 genes) and 3dpi (184 genes). RT-qPCR validation of selected ISGs showed that ISG15, LAMP3, SAMHD1, RASGRP3, and C19ORF66 were induced in infected FRPE cells at both 2 and 4 dpi (Figure 2C). Moreover, infected cells had induced expression of pro-inflammatory genes TNFα, IL-6, and ICAM1 (Figure 2C). Mouse studies have shown that ZIKV directly inhibits STAT2 during infection (Grant *et al*., 2016). In human FRPE cells, key JAK-STAT pathway transcription factors were upregulated during infection, including IFNAR1, IFNAR2, and STAT1 (Figure 2 and Tables S2-S3). Additionally, western blot analysis showed induction of total STAT1 protein in ZIKV infected cells at 4 dpi (Figure 2D). Analysis of gene ontology categories and functional pathways showed the activation of various genes involved in the inflammasome pathway (NLRP1, NLRP3, ASC, AIM2, and NLRC4), IL-17 signaling pathway, and the VEGFA signaling pathway, which is part of tissue healing and repair processes (Figure S3 and Tables S2-S3). Infected FRPE cells showed increased levels of VEGFA as compared to mock cells (Figure 2C). Proteomics analysis identified 492 deregulated proteins (*p*-value of less than 0.05) involved in diverse cellular processes, which included cell growth and proliferation, cell survival, metabolism, transcription, and protein synthesis (Table S4). The top survival pathways identified in both transcriptomics and proteomics analyses were PI3/AKT and ERK/MAPK (Figure 3A). Moreover, several proteins involved in cell survival (DDX5 and PHB), phagocytosis (TXNDC5), particle internalization (RPSA), and oxidative stress response (PRDX2 and TCP1) were differentially regulated (Figure 3B). Proteins involved in controlling mitochondrial membrane potential (CLIC1, CLIC4, and LDH4) were deregulated at both 2 and 4 dpi (Figure 3C). The Rac signaling pathway was one of many top hits in the proteomics common pathway analysis (Figure 3A). Overall, our results indicate that fetal retinal pigment epithelial cells respond to ZIKV infection by activating innate immune, inflammatory, and stress response pathways.

**Figure 2.**
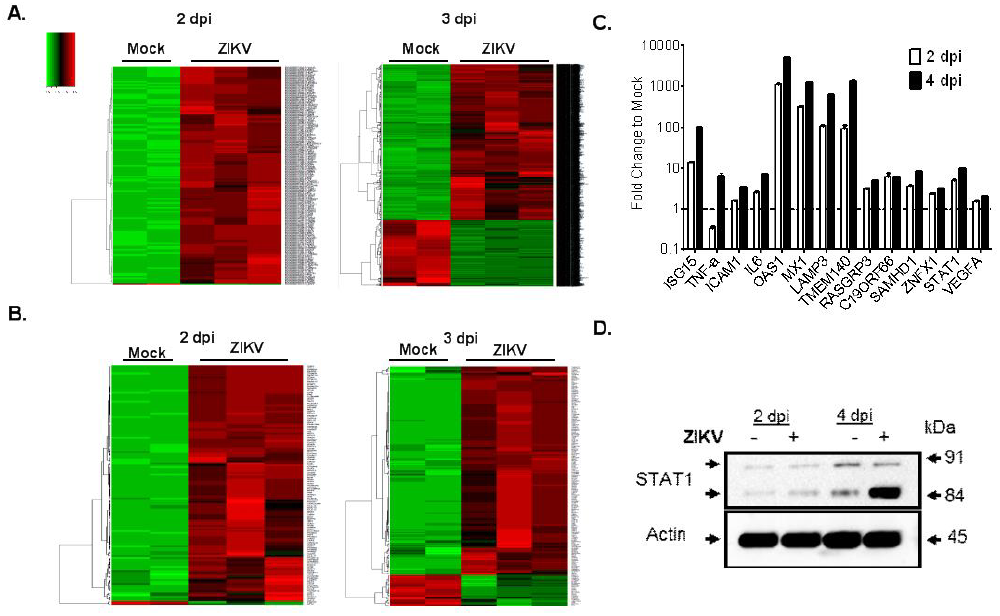
ZIKV dysregulates host cellular processes and activates innate immune and inflammatory responses in fetal RPE cells. (A) Heat map of differentially expressed genes at 2 dpi (2,040 genes) and 3 dpi (7,951 genes) [upregulated (red); downregulated (red)]. 2-way supervised Hierarchical Clustering was performed using unique genes identified for the combined dataset. (B) Heat maps show the differential expression of known canonical ISGs in mock and ZIKV infected cells at 2 dpi (128 genes) and 3 dpi (184 genes). (C) RT-qPCR validation of selected inflammatory and IFN-stimulated genes from (B) in ZIKV infected FRPE cells at 2 and 4 dpi. The fold change to mock is shown. The horizontal dash line represents the mock value at 1 where the fold change was taken for the specified time points. (D) Western blot shows induction of STAT1 after ZIKV infection at 2 and 4 dpi. Note that higher levels of STAT1 beta subunit (84 kDa) are present in infected cells. Actin was included as a loading control. (D) Key pathways identified during ZIKV infection of FRPE cells through comprehensive transcriptomics and proteomics analyses. ZIKV infections were done in biological triplicates for each time point. The uninfected controls were done in biological quadruplicates and pooled in duplicates for downstream analyses.

**Figure 3.**
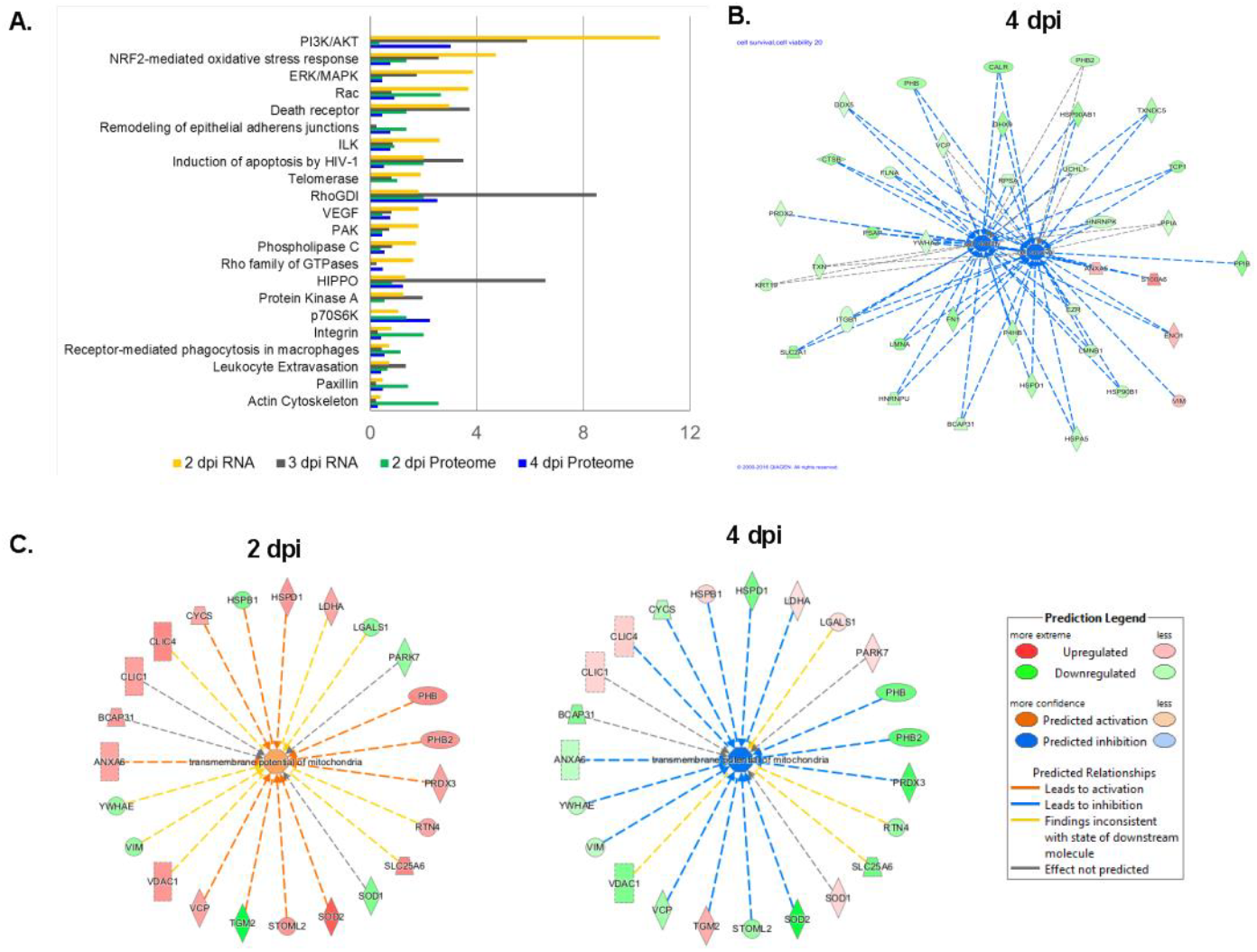
Proteomics analysis of human fetal RPE cells infected with ZIKV. (A) Graph shows key deregulated signaling pathways identified in FRPE cells through comprehensive proteomics and transcriptomics proteomics analyses during ZIKV infection. Interaction networks of dysregulated factors involved in cell survival (B) and mitochondrial transmembrane potential (C) during ZIKV infection are presented. [Upregulated (red); downregulated (green)]. For proteomics experiments, biological quadruplicates were performed and then samples were pooled in duplicate for mass spectrometry.

### Drug treatment to reduce ZIKV-mediated ocular cell death

We then performed a small screen using selected compounds in Vero cells infected with ZIKV (Figure 4A), which have been previously shown to have anti-flaviviral activity. From this screen, we identified 6-Azauridine (6-AZA) (Figure 4B), a nucleoside analogue, as a potent inhibitor of ZIKV. Subsequently, we tested 6-AZA to test its antiviral activity against ZIKV in the context of the biologically relevant fetal RPE cells. The drug compound cyclopentenyl cytosine (CPEC) has also been shown to have a broad spectrum anti-viral effect on West Nile virus (Morrey et al., 2002), which did not inhibit ZIKV in Vero cells (Figure 4A). To determine the effects of each compound on ZIKV-mediated apoptotic cell death, we measured caspase 3/7 activity after ZIKV infection and drug treatment (1 μM, 5 μM, and 10 μM) at 72 hpi. There was a significant decrease in caspase 3/7 activity upon treatment with 6-AZA at the 5 μM and 10 μM concentrations (Figure 4C), which correlated with the significant decrease in ZIKV infection (IC_50_ of 5 μM) (Figure 4D). CPEC had similar caspase 3/7 activity as the ZIKV-only infected group (vehicle) (Figure 4C). Immunostaining confirmed the anti-viral activity of 6-Azauridine, which showed significantly reduced viral foci formation (5 and 10 μM) as compared to the vehicle control (Figure 4E). Whereas, there was no change observed in ZIKV replication in the CPEC treated groups, which had a similar level of infected cell foci as the vehicle control (Figure 4E). Overall, the nucleoside analogue, 6-AZA, had an anti-Zika viral effect and reduced the apoptotic cellular response in fetal RPE cells.

**Figure 4.**
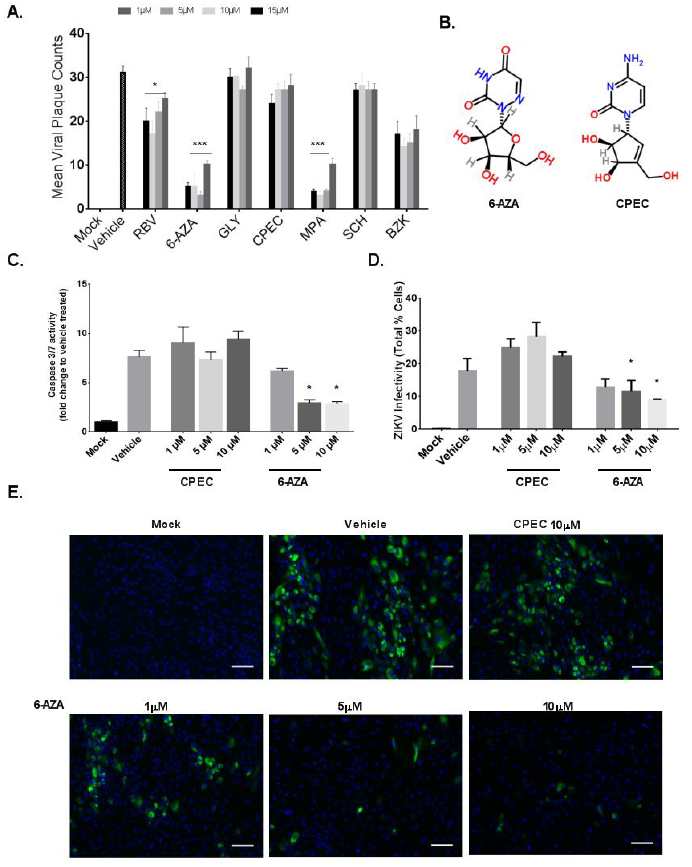
6-Azauridine inhibits ZIKV replication in FRPE cells. (A) Graph depicting mean Zika viral plaque counts in an antiviral chemical compound screen in Vero cells. Note: RBV (ribavirin); 6-Aza (6-azauridine); GLY (Glycyrrhizin); CPEC (cyclopentenyl cytosine); MPA (mycophenolic acid); SCH (sodium cholate hydrate); BZK (benzalkonium chloride). (B) Chemical compound structure of 6-AZA and CPEC. (C) Caspase 3/7 activity measured for mock (control), ZIKV infected cells (Vehicle treated) (MOI of 1), and infected, but treated cells with CPEC and 6-Azauridine at 1, 5, and 10 μM concentrations. (D) Bar graphs depicting the percentage of ZIKV infected cells (% cells) in a well of a 96-well plate as detected by ZIKV Env antibody in each of the treated and control groups. (E) Immunocytochemistry images show mock, vehicle, and ZIKV infected and drug treated cells. Treatment with 6-AZA shows a dose dependent anti-viral effect in FRPE cells. Scale bar: 100 μm. Student t-test was performed for statistical significance if the p value was p<0.05 (*); p<0.001 (**); and p<0.0001 (***). Representative data from three independent experiments are shown.

### iPS-derived retinal stem cells (iRSCs) are susceptible to ZIKV infection

We then evaluated the susceptibility of iRSCs, also known as eye field stem cells (EFSCs) (Zhao and Afshari, 2016), to ZIKV infection. iRSCs represent the multi-potent retinal stem/progenitor cell population present in the first trimester of the developing human eye. iRSCs are able to differentiate into RPE cells, photoreceptors, retinal ganglion cells, and corneal endothelial cells. Human retinal stem cells were differentiated from iPSCs under chemically defined conditions, which were originally derived from BJ human fibroblast cells (ATCC). The BJ-iPSCs (passage 7) were directionally induced to form retinal stem cells as shown in Figure 5A (Zhao and Afshari, 2016). The expression of retinal stem cell transcription factor genes PAX6 and LHX2 were confirmed in the generated iRSCs (Figure 5B). The differentiated cells were infected with ZIKV at an MOI of 0.1. iRSCs were susceptible to ZIKV infection as shown by the presence of viral Env protein (green) in the cytoplasm of ZIKV-infected iRSCs at 2 and 4 dpi (Figure 5C). iRSCs showed increased infectivity at 4 dpi, as reflected by ZIKV genome copies (Figure 5D). iRSC viability was affected by 4 dpi with a 35% reduction in infected cells as compared to mock cells (Figure 5E). Although there was an increase in Caspase-3/7 activity by 4 dpi, it was not statistically significant. In comparison to ZIKV infected FRPE cells, iRSCs are less susceptible to ZIKV infection, as iRSCs had a 4 log lower level of viral genome copies to that of FRPE cells at 2 dpi under similar infection conditions (Figures 1C and 5D). The iRSCs showed lowered expression of the selected flaviviral entry receptors AXL, heat shock proteins (HSP90AA1, and HSPA5), and integrins (ITGA3, and ITGA5) when compared to FRPE cells (Figure 5E and Figure S1B). These receptors may partially explain the higher susceptibility of FRPE cells to ZIKV infection. It is possible that other intrinsic factors could contribute to the differential susceptibility. Overall, ZIKV-infected iRSCs showed a delay in viral infection, with a significant decrease in cell viability, and increased ZIKV genome copy number by 4 dpi.

**Figure 5.**
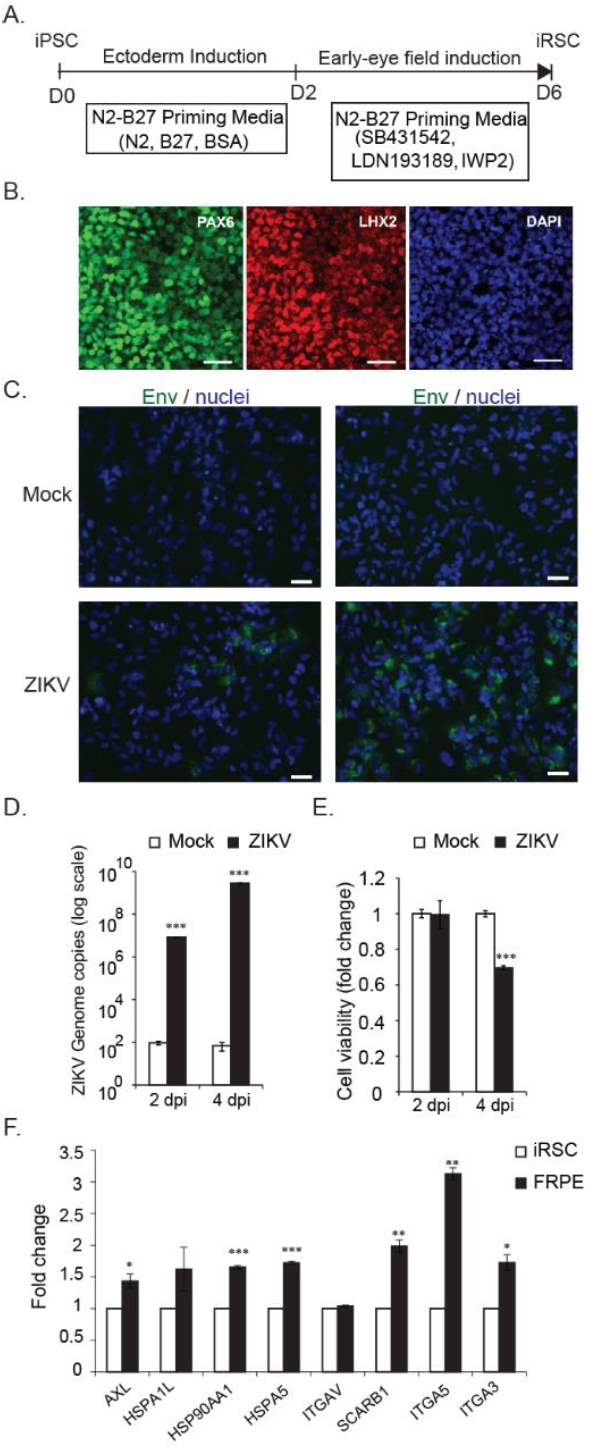
iPSC-derived retinal stem cells (iRSCs) for ZIKV infection. (A) Schematic diagram shows the experimental outline used for the differentiation of iPSCs into iRSCs. (B) Differentiated iRSCs show the expression of early eye stem cell transcription factors PAX6 and LHX2. Nuclei are stained with DAPI. Scale bar: 30 μm. (C) Immunocytochemistry images of mock and ZIKV-infected iRSCs at 2 dpi (left panels) and 4 dpi (right panels). The presence of viral Env protein (green) is verified by immunostaining. Nuclei are stained with DAPI (blue). Scale bar: 25 μm. (D) Graph shows the quantification of ZIKV genome copies in mock and ZIKV infected iRSCs at 2 and 4 dpi. (E) Viability of iRSCs mock and ZIKV infected 2 and 4 dpi. The fold change to mock is shown. (F) Graph depicts the expression of flaviviral entry receptors on FRPE cells as compared to iRSCs. Student t-test was performed for statistical significance if the p value was p<0.05 (*); p<0.001 (**); and p<0.0001 (***). Representative data from three independent experiments are shown.

## Discussion

The severity of ocular cell injury caused by ZIKV may contribute to the outcome of congenital eye disease (de Paula Freitas et al., 2016; Miner et al., 2016). Our studies provide insight on the initial cellular response to ZIKV infection by examining cells that comprise different stages of eye development including iRSCs, which are multipotent progenitor cells that are equivalent to cells found in the first trimester of development, and human fetal RPE cells, which constitute the cells of the retina at the second-trimester (∼20 weeks) (Reh and Fischer, 2006; Völkner et al., 2016). We first evaluated the permissiveness of FRPE cells to ZIKV infection. We found that FRPE cells are highly susceptible to ZIKV infection, further leading to increased apoptosis and reduced cell viability. Transcriptomics analysis of infected FRPE cells showed a type I interferon signature indicative of innate immune pathway activation. Furthermore, the infected FRPE cells responded by activation of inflammasome and IL-17 signaling pathways. These pathways can amplify the inflammatory state by recruiting first-line of defense cells, such as neutrophils and macrophages, to the infected site *in vivo*. Furthermore, proteomics analyses revealed the dysregulation of factors involved in cell survival, such as the PI3K/AKT signaling pathway.

During development, retinal stem cells are precursor progenitor cells to RPE cells, retinal ganglion cells, corneal endothelial cells, and photoreceptors (Zhao and Afshari, 2016; Parameswaran et al., 2010; Osakada et al., 2008, Nakano et al., 2013, Zhong et al., 2014). For this reason, iRSCs are a useful *in vitro* model system to decipher the pathobiology of ZIKV eye infection at the early stages of ocular development. In this study, we showed that retinal stem cell is susceptible to ZIKV infection, but provides a lower degree of infection as compared to its downstream lineage FRPE cells. This suggests that ZIKV has differential permissivity to the state of cellular maturity of the cells. Furthermore, analysis of flaviviral entry receptor expression showed that FRPEs had higher levels of AXL, heat shock proteins, and integrins than iRSCs, which may explain differential susceptibility of these cell types. It is also possible that iRSCs may lack other pertinent cellular factors critical for robust ZIKV replication. ZIKV infection may impact the development of multiple cell fates, further leading to the generation of aberrant non-functional cells and/or malformations of the eye in the developing fetus. Moreover, optic cup cells, including photoreceptors are derived from retinal stem cells and if the RSCs are damaged by ZIKV during the first trimester, then abnormal development of the optic cup, will ensue. Further studies using a retinal organoid model system (Völkner et al., 2016) may provide insight on how retinal, choroidal, and different optic cells are affected during ZIKV infection. RPE cells function as a support cell type providing nutrients, growth factors, and gas exchange to photoreceptors and other retinal cells. Death of RPE cells would indirectly affect the health of photoreceptors. Furthermore, ZIKV infection may lead to changes in the retinal architecture by altering RPE tight junctions and choroidal vascular permeability.

Recent studies show that ZIKV can halt the cell cycle and induce apoptotic cell death in human neuroprogenitors and brain organoids (Dang et al., 2016; Garcez et al., 2016; Tang et al., 2016). In our studies, ZIKV induced caspase 3/7 activity in FRPE cells. These results suggest that the tissue injury caused by apoptotic cell death could be a major contributing factor to the eye pathology observed in fetuses and affected infants. Viral particles released by apoptotic bodies can be taken up by neighboring cells and macrophages *in vivo*, in turn facilitating viral spread (Roulston et al., 1999). There is also evidence for active ZIKV infection of macrophages in the placenta of pregnant mothers, which in turn infect the developing fetus (Jurado et al., 2016). In addition, proteomics analyses of infected FRPE cells showed activation of proteins controlling mitochondrial membrane potential. This increased mitochondrial potential could be due to the infected cell undergoing active apoptotic processes. The tissue damage caused by other flaviviruses can also be attributed to the activation of the programmed cell death pathway. For example, platelets from Dengue-infected patients show characteristic apoptotic features, such as the induction of caspase -3 and -9, as well as mitochondrial polarization (Hottz et al., 2013).

A recent study describing the humoral response during ZIKV infection of pregnant women with neonatal microcephaly, showed that the amniotic fluid had increased levels of VEGF, pro-inflammatory cytokines (IL-8 and IL-6), and chemokines (IP-10 and MCP-1), which may contribute to other fetal developmental abnormalities (Ornelas et al., 2016). Consistent with this study, we observed that ZIKV induces the expression of VEGF and various chemokines (CXCL1 and CCL5) in fetal RPE cells. RPE cells require VEGF for the health of the endothelium and underlying blood vessels (Ford et al., 2011; Nagineni et al., 2012). VEGF binds to the receptor Flt1 and KDR on endothelial cells and can initiate different intracellular signaling pathways, including PI3/AKT, FAK, PKC, and MAPK pathways to promote cell survival and angiogenesis (Eichmann and Simons, 2012). VEGF has also been shown to be involved in the pathogenesis of an important degenerative eye disease. In wet form of age-related macular degeneration (AMD) disease, overproduction of VEGF leads to abnormal growth of choriocapillaris (Kovach et al., 2012). Congenital eye lesions caused by ZIKV may resemble macular degeneration (de Paula Freitas et al., 2016). *In vivo* studies may provide insight on the role of VEGF during ZIKV-mediated eye disease.

To investigate the underlying molecular mechanisms of congenital ZIKV-mediated eye disease, a mouse pregnancy model that utilizes intraocular injection of ZIKV in the fetus (E8.5-9 stage) can be developed. Moreover, it will be important to investigate the Zika virulence factors that contribute to congenital eye injury and disease pathogenesis. Detailed analysis of the immunologic features of infected ocular cells *in vivo* may enable the development of immunotherapeutic strategies against ZIKV.

Currently, there are no approved antiviral agents to treat and prevent ZIKV-mediated disease. In this study, we tested drug compounds previously shown to have antiviral effects on different flaviviral pathogens, including Dengue, Yellow Fever, and West Nile viruses (Crance et al., 2003). Our *in vitro* studies show that treatment with 6-AZA blocked virus production and apoptosis in FRPE cells immediately after ZIKV infection. 6-AZA was recently shown to have anti-ZIKV activity (Pascoalino et al., 2016; Adcock et al., 2017). In general, nucleoside analogues get incorporated into the newly synthesized viral genome, further causing deleterious mutations. Furthermore, the ZIKV polymerase NS5 may be a target for such nucleoside analogues. Although vaccines are the best choice for preventing ZIKV infection in pregnant women, antiviral drugs can be used as a post-exposure therapeutic option.

In summary, this study provides a snapshot into ZIKV infection at the first and second-trimester of ocular development using biologically relevant *in vitro* ocular cell model systems. Our study presents a comprehensive assessment of pathways affected in fetal ocular cells by transcriptomics and proteomics analyses. This data set will serve as a resource to the scientific community to further dissect the pathobiology of ZIKV-mediated eye and brain diseases.

## Experimental Procedures

### Cells

Human fetal retinal pigment epithelial (FRPE) cells (kindly provided by Dr. Guoping Fan at UCLA) were grown in DMEM high glucose medium supplemented with 10% fetal bovine serum and 1% penicillin/streptomycin. FRPEs were originally isolated from fetal retina (∼20-week-old). The cells were incubated at 37^°^C supplemented with 5% CO_2_ and passaged when 90% confluency was reached at approximately every third day using 0.05% Trypsin with 0.53 mM EDTA (Corning, USA).

Induced pluripotent stem cells (iPSCs)-derived retinal stem cells (iRSCs): The iPSC line was derived from BJ (ATCC^®^ CRL-2522) human fibroblast cells (American Type Culture Collection, Manassas, VA, USA). At passage 7, iPSCs were induced into retinal stem cells (ABC010) by chemically defined conditions, as previously described (Zhao and Afshari, 2016., 2016). In brief, induced pluripotent stem cell-derived retinal stem cells (iRSCs) were grown in DMEM/F12 media supplemented with 1% N2, 2% B27 without vitamin A, 7.5% BSA Fraction V, 1% GlutaMAX, 1% MEM non-essential amino acids, 1% of a proprietary cocktail (provided by Alpine Biotech LLC) for retinal stem cells, and 300 μM Ascorbic acid on matrigel coated flasks. Dissociation of the cells was achieved by Accutase (Life Technologies, USA). The expression of retinal stem cell markers was verified.

### Zika virus

The PRVABC59 (GenBank accession number KU501215) Zika virus strain of Asian genotype was used for the infection of human retinal pigment epithelial (FRPE) cells and iRSCs. PRVABC59 was acquired from the Centers for Disease Control and Prevention (CDC), USA. Working viral stock for the specified experiments was generated by subjecting the original ZIKV strain (Passage=3) to two additional passages in Vero cells. An established viral plaque assay was utilized to measure viral titer, as previously described (Contreras and Arumugaswami, 2016).

### Zika viral infection

For ZIKV infection, FRPE and iRSCs were seeded in a 24-well plate with a cell density of ∼2.5 × 10^4^ cells/well. After 24-hours, ZIKV inoculum, with a multiplicity of infection (MOI) of 0.1, was formulated using the base media specified for each cell type. A total of 200 μL of viral inoculum was added to each well and the plates were incubated at 37 °C with 5% CO_2_ for 2-4 hrs. After the incubation period, the base media was replaced for each cell type with their respective complete media at a volume of 1 mL per well. As for the uninfected (mock) group, each cell type received the specified cell-growth media that was concurrently used to prepare the viral inoculum, as described above. Each cell type included a mock infected control for the specified timepoints of 2 and 3 days post infection, for relevant comparison. At the end of each timepoint, cell lysates and RNA were harvested for downstream analyses.

### Caspase 3/7 assay to measure apoptosis

Caspase-Glo 3/7 Assay (Promega, USA) was performed as per the manufacturer’s protocol. At the indicated time points, ZIKV infected and mock infected FRPE cells and iRSCs were incubated with the pro-luminescent caspase-3/7 substrate for 1 hour at room temperature. Subsequently, 100 μL of lysate was transferred to a white 96-well microtiter plate for reading the luminescence signal using a luminometer (Glomax Microplate Luminometer, Promega).

### Proteomic sample preparations and Liquid chromatography Mass Spectrometry (LC-MS/MS) analysis

For proteomics experiments, biological quadruplicates were performed and subsequently pooled into duplicates for sample analysis using mass spectrometry. After FRPE cells were lysed in 80 mM Tris-HCl, 4% SDS, 100 mM DTT at pH 7.4, 30 μg of protein was quantified for each sample and further digested using the FASP Protein Digestion Kit (Expedeon, San Diego, CA, USA) per manufacturer’s instructions.

Liquid chromatography tandem mass spectrometry (LC-MS/MS) system was set up on a Transcend II LX-2 HTLC system connected to an Q exactiveplus equipped with an HESI ion source (QE, Thermo Scientific). A LC-MS/MS setup was used to separate the tryptic digested peptides with mobile phases A (0.1% formic acid) and B (0.1% formic acid in 100% acetonitrile). The tryptic peptide mixture (20 μg) was loaded onto a large-inner-diameter LC column (Onyx™ Monolithic C18, LC Column 50 × 4.6 mm, 130 A) and desalting for 3 min with 2% mobile phase B at a flow rate of 200 μL/min. A series of gradients (flow rate, 200 μL/min) was used to elute the trapped peptides for separation and then they were further applied to a Mass Spectrometer. The complex peptide mixture was separated using a 120-min linear gradient, which consisted of the following steps: 2% B for 5 min, 3% to 35% B in 100 min, 35% to 95% B in 20 min, then maintaining isocratic conditions at 95% B for 10 min, and finally reverting back to 2% B for 20 min of re-equilibration.

The mass spectrometer QE was operated in positive ion mode with the data-dependent acquisition strategy. Capillary temperature was set to 320°C and spray voltage was set to 3.5 kV with sheath gas of 40, auxiliary gas of 20, S lens voltage was 55 volts and hearer temperature was 350°C. One scan cycle included an MS1 scan (*m/z* 400–900) acquired at a resolution (full width at half-maximum) of 70,000 with an AGC target of 3 × 10^6^ ions over a maximum time of 100 ms, followed by ten MS2 scans with resolution of 17,500 at in source collision-induced dissociation activation mode to fragment the ten most abundant precursors found in the MS1 spectrum with a target setting of 1 × 10^5^ ions, an accumulation time of 50 ms, and an isolation window of 4 Da (m/z). Normalized HCD collision energy was set to 27%, and one microscan was acquired for each spectrum. The dynamic exclusion was enabled with the following settings: repeat count, 1; repeat duration, 30 s; exclusion list size, 500; exclusion duration, 20 s. In this study, two respective cell lysates of control groups (Mock, day 2 and day 4) and experimental groups (Infected, day 2 and day 4) were analyzed sequentially.

### Proteomic bioinformatics

To identify proteins in mock (control) and ZIKV infected FRPE cells, mass spectroscopy data was processed by Trans-proteomic Pipeline software (Deutsch et al., 2010; Keller et al., 2005) and Skyline daily v.3.5 software (MacLean et al., 2010). Fold changes between infected and mock samples were generated using intensity values from the MPPReport plugin in Skyline daily. To identify cellular processes that are affected during ZIKV infection, the list of proteins from infected vs. mock samples at 2 and 4 dpi were input into Ingenuity Pathway Analysis (Qiagen, Redwood City, CA, USA) and network diagrams were generated.

### RNA library preparation and sequencing

For RNA sequencing, FRPE cells were infected with ZIKV at an MOI of 1 and cell lysates were harvested at 2 and 3 dpi. ZIKV infections were done in biological triplicates for each time point. The uninfected controls were done in biological quadruplicates and pooled in duplicates for downstream analyses. Total RNA was isolated from the cells using an RNeasy Mini Kit (QIAGEN, USA). RNase-free DNase treatment was performed on the column to remove residual DNA. Library preparation and RNA sequencing analysis were done by Cedars-Sinai Genomics Core. Samples were assessed for concentration and quality using the Thermo Scientific NanoDrop 8000 Spectrophotometer (Waltham, MA). RNA sequencing libraries were constructed using Illumina TruSeq RNA Sample Preparation Kit v2 (Illumina, San Diego, CA), as per manufacturer’s instructions. Briefly, total RNA, with RNA integrity (RIN) scores of 9 or better (Agilent Bioanalyzer RNA 6000 Nano kit) were used for RNA library preparation. The poly (A)+ RNA was then purified from one microgram of total RNA from each sample using oligo-dT attached magnetic beads with two rounds of purification.

Subsequently, the poly (A)+ RNA was fragmented and primed for cDNA synthesis per manufacturer’s recommendations. RNA adapters and barcodes were ligated to cDNA to allow for clonal amplification and multiplexing. Sequencing was done on an Illumina NextSeq 500 using 75 bp single-end sequencing kit to yield an average read depth of 28 million reads per sample with a minimum number of reads for a sample of at least 16 million reads. The sequencing data was deposited to the Gene Expression Omnibus (GEO) database system with the accession number GSE83900.

### Analysis of RNA sequencing data

Raw reads obtained from RNA-sequencing were aligned to the human genome using STAR (version 2.5) (Langmead et al., 2009) with a custom human GRCh38 transcriptome reference downloaded from http://www.gencodegenes.org, which contains all protein-coding and non-coding RNA genes based on human GENCODE version 23 annotation.

Expression counts for each gene in all the samples were normalized by a modified trimmed mean of the M-values normalization method. The unsupervised principal component analysis (PCA) was performed with DESeq2 Bioconductor package version 1.10.1 in R version 3.2.2 (Love et al., 2014). Each gene was fitted into a negative binomial generalized linear model. The Wald test was applied to assess the differential expressions between two sample groups by DESeq2. The Benjamini and Hochberg procedure (Benjamini and Hochberg, 1995) was applied to adjust for multiple hypothesis testing and differentially expressed gene candidates were selected with a false discovery rate of less than 0.10. For visualization of coordinated gene expression in the samples, a hierarchical clustering with the Pearson correlation distance matrix was performed with differentially expressed gene candidates using the Bioconductor gplots package (version 2.14.2) in R. Significantly expressed genes were assessed for pathway enrichment using DAVID release 6.7 (https://david.ncifcrf.gov/) and Ingenuity Pathway Analysis (http://www.ingenuity.com/products/ipa) (QIAGEN, Redwood City). The significantly enriched canonical pathways were defined as having a q-value of <0.01.

### Reverse Transcription-Quantitative PCR Analysis

Total RNA was extracted from mock and ZIKV-infected FRPE and iRSCs at the designated time points using an RNeasy Mini Kit (QIAGEN). After treatment with RNase-free DNase, 1 μg of RNA was reverse-transcribed into cDNA using random hexamer primers and the SuperScript III Reverse Transcriptase Kit (Life Technologies), as recommended by the manufacturer. The following conditions were used for cDNA amplification: 65°C for 5 min and 4°C for 1 min, followed by 55°C for 60 min and 72°C for 15 min. Quantitative real-time PCR was carried out using Platinum SYBR Green qPCR SuperMix-UDG with ROX Kit (Life Technologies) by the QuantStudio™ 12K Flex Real-Time PCR System (Life Technologies). The relative concentration of each transcript was calculated using 2^−ΔCT^ method using Glyceraldehyde 3-phosphate dehydrogenase (GAPDH) threshold cycle (C_T_) values for normalization. The qPCR primer pairs for the mRNA transcript targets are provided in Table S1. The following conditions were used for transcript amplification: 50°C for 2 min and 95°C for 2 min, followed by 40 cycles of 95°C for 15 sec and 60°C for 1 min.

### Immunocytochemistry

Mock and infected FRPE cells and iRSCs were fixed with ≥99.8% methanol (Sigma-Aldrich, St. Louis, MO) and incubated at −20°C for 30 minutes, then washed three times with 1X DPBS (Dulbecco’s Phosphate-Buffered Saline (DPBS) (Corning Incorporated., Manassas, VA). The cells were permeabilized with 3% Bovine Serum Albumin (BSA) (Sigma Life Sciences, St. Louis, MO) and 0.1% Triton-X 100 in 1X DPBS. For ZIKV immunostaining, fixed and permeabilized cells were incubated with mouse monoclonal antibody against the Flavivirus Envelope protein (Absolute Antibody Ltd.) at a 1:200 dilution for up to 6 hrs or overnight incubation at 4^°^C. The secondary antibodies, goat anti-mouse polyclonal antibody (Alexa Fluor 488) or goat anti-mouse polyclonal antibody (Alexa Fluor 594) (Life Technologies, USA) were added at 1:1000 dilutions and incubated for 1 hr at room temperature. For staining of RPE markers, fixed cells were incubated overnight at 4°C with anti-bestrophin (BEST-1) mouse monoclonal primary antibody (EMD Millipore, Billerica, MA), anti-retinal pigment epithelium 65 (RPE 65) monoclonal primary antibody (EMD Millipore, Billerica, MA), and anti-tight junction protein 1 (zona occludens 1) (TJP1/ZO-1) rabbit polyclonal antibody (Life Technologies, Grand Island, NY), all at a dilution of 1:200. The secondary antibodies, goat anti-mouse polyclonal antibody (Alexa Fluor 594) (Life Technologies, Grand Island, NY) and goat anti-rabbit polyclonal antibody (Alexa Fluor 568) (Life Technologies, Grand Island, NY) were added at a dilution of 1:200, and were incubated for an hour at room temperature. Cells were washed 3 times with 1X DPBS and left on the rocker for 5 minutes for each wash and in between antibody changes. Nuclei was stained with DAPI (4’,6-Diamidino-2-Phenylindole, Dihydrochloride) (Life Technologies., Grand Island, NY) at a dilution of 1:1000.

### Antiviral compound analysis

A compound screen in a dose-response was performed in Vero cells. The cells were seeded at a cell density of 1×10^4^ cells per well in 96-well plates. After 16 hours of plating, the cells were infected with ZIKV (MOI of 1). After 4 hpi, the cells were treated with test compounds, ribavirin (RBV), 6-azauridine (6-AZA), glycyrrhizin (GLY), cyclopentenyl cytosine (CPEC), mycophenolic acid (MPA), sodium cholate hydrate (SCH), benzalkonium chloride (BZK) and curcumin (CRM) at the following concentrations of 1 μM, 5 μM, and 10 μM in triplicates. All compounds were acquired from Sigma Aldrich unless otherwise stated. Cells were infected and treated for 72 hours, at which point viral plaques were counted. Similarly, FRPE cells were seeded at a cell density of 1×10^4^ cells per well in 96-well plates. After 16 hours of plating, the cells were infected with ZIKV (MOI of 1) and immediately following infection, test compounds were added at the following concentrations of 1 μM, 5 μM and 10 μM in triplicates. Cells were infected and treated for 72 hours, at which point cell viability and apoptosis was measured, and the antiviral activity was measured. The mock, vehicle (ZIKV), and infected plus compound treated samples were immunostained, as mentioned above, and were analyzed using ImageXpressMICRO (Molecular Devices, USA) with multi-wavelength cell scoring macro software. For each well, the number of nuclei (DAPI) and cell count was determined for nine different locations within the well. Data acquisition was based on DAPI-normalization.

### Western Blot analysis

Cell lysates were resolved by SDS-PAGE using 4-15% pre-cast gradient gels (Bio-Rad., Hercules, CA) before using the Trans-Blot turbo transfer system (Bio-Rad., Hercules, CA) to transfer to a 0.2 μm PVDF membrane. Subsequently, the membranes were blocked with 5% skim milk and 0.2% Tween-20 in PBS at room temperature for 1 hour. The membrane was then probed with rabbit monoclonal antibodies STAT1 and beta-actin (Cell Signaling Technology, Danvers, MA). Stabilized Peroxidase Conjugated Goat anti-rabbit IgG (H+L) secondary antibody (Thermo Scientific, Grand Island, NY) was added and detected by chemiluminescence (SuperSignal West Pico Chemiluminescent Substrate kit, Thermo Scientific, Grand Island, NY).

### Statistical analysis

Three independent biological replicates were carried out for each of the represented experiments in the study. The error bars in the graph reflect the standard deviation. P-values were determined by a two-tailed student’s t-test and significance was reported if the p value was p<0.05 (*); p<0.001 (**); and p<0.0001 (***). For reproducibility, experiments were repeated two or more times.

## Author Contributions

D.C: designed and executed experiments, analyzed and interpreted data, and wrote the manuscript. M.K: performed experiments and analyzed data. L.E.M: performed experiments, analyzed data and wrote the manuscript. V.G. and J.T: analyzed and interpreted RNA sequencing data. Z.C: performed proteomics experiments and analyzed data. Y.W. and J.Z: provided reagents and performed iRSC induction experiments. S.W: provided reagents and analyzed and interpreted data. V.A: designed experiments, analyzed and interpreted data, and wrote the manuscript.

## Acknowledgements

We thank Dr. Gouping Fan (UCLA, USA) for kindly providing the human fetal retinal pigment epithelial cells; Dr. Aaron Brault and Dr. Brandy Russell of the Centers for Disease Control and Prevention (CDC, USA) for providing the Zika viral strain PRVABC59; and Gustavo Garcia Jr. for technical support and editing the manuscript.

